# Heat-responsive dynamic shifts in alternative splicing of the coral *Acropora cervicornis*

**DOI:** 10.1101/2025.01.21.634199

**Authors:** Kathryn H. Stankiewicz, Jacob J. Valenzuela, Serdar Turkarslan, Wei-Ju Wu, Kelly Gomez-Campo, Nicolas S. Locatelli, Trinity L. Conn, Veronica Z. Radice, Katherine E. Parker, Rachel Alderdice, Line K. Bay, Christian R. Voolstra, Daniel J. Barshis, Iliana B. Baums, Nitin S. Baliga

## Abstract

Climate change has caused drastic declines in corals. As sessile organisms, response to shifting environmental conditions may include changes in gene expression, epigenetic modifications, or the microbiome, but as of yet, a common mechanism of stress response, alternative splicing (AS), has been underexplored in corals. Using short-term acute thermal stress assays, we investigated patterns of AS in the scleractinian coral *Acropora cervicornis* during response to and a subsequent overnight recovery phase from low (33℃), medium (35℃), and high (37℃) levels of heat stress. We find that 40% of the genomic gene set is subject to AS. Our findings demonstrate conserved and dynamic shifts in splicing profiles during the heat treatment and subsequent recovery phase. AS increased in response to heat stress and was primarily dominated by intron retention in specific classes of transcripts, including those related to splicing regulation itself. While AS returned to baseline levels post-exposure to low heat, AS persisted even after reprieve from higher levels of heat stress. Partial overlap of AS transcripts with differentially expressed genes suggests that AS may represent a distinct and previously underappreciated regulatory mechanism for thermal stress response in corals.

## Introduction

Scleractinian corals are early branching metazoans responsible for building the three-dimensional structure of reef ecosystems which house 32%^1^ of ocean biodiversity, and thus, critical pillars of healthy marine environments and coastal communities^2,3^. Given their ecological, environmental, and social importance, it is cause for great concern that coral reefs are declining substantially due to climate change^4,5^. The breakdown of the partnership between the coral host and its photosynthesizing algal endosymbiont (family Symbiodiniaceae)^6^, known as ‘coral bleaching’, is triggered by stress and impedes symbiotic nutrient cycling often leading to subsequent mortality of the coral host^7,8^. Widespread bleaching of coral populations across entire reef tracts (‘mass bleaching events’) due to temperature stress–especially heat related–have become more frequent in recent years and many coral species are now critically endangered^9,10^. Consequently, it is important to gain an understanding of the molecular mechanisms driving disparate outcomes following heat stress across and within reefs.

Much effort in coral conservation has gone towards understanding the mechanisms behind the coral-algal (‘holobiont’) response to stress. In particular, many studies have examined the transcriptional response of the holobiont to stress^11–13^. This has uncovered the role of particular genes and pathways in the environmental stress response, such as components of the apoptotic signaling cascade, lectins, and genes related to nucleic acid repair, growth arrest, oxidative stress, and heat shock proteins^14–17^. Differences in thermal sensitivities have also been described between corals based on phylogenetic lineage and species^18^, microbiome^13,19^, population^13^, location^20^, as well as the associated symbiont species^21,22^. Innate thermal stress sensitivity of the host and symbiont acts in concert with variation in environmental condition^23^. In recent years, there is also evidence that epigenetic mechanisms may play a role in acclimatization^24–27^. However, to our knowledge, no study to date has investigated the impact of thermal stress in corals at the level of mRNA splicing - an established mechanism of stress adaptation used by plants^28^.

Eukaryotic alternative splicing (AS) of pre-mRNA produces multiple transcript isoforms from a single gene, and thus, can increase proteome diversity and complexity without requiring changes to the underlying genomic sequence^29^. Under constitutive splicing, introns are excised and exons are ligated to produce mature mRNA transcripts—a process mediated by the spliceosome, a macromolecule of ribonucleic proteins^30–34^. Alternative splicing may result in an alternative 3’ or 5’ splice site, multiple or single skipped exons, mutually exclusive exons, or intron retention^35,36^. AS has been shown to be universal among plants and animals and implicated in various cellular responses, e.g., human diseases^37^, eukaryotic tissue development and differentiation^38^, and response to abiotic stress^28^.

Here, we characterize the global AS landscape by elucidating the splicing response of a Caribbean coral, *Acropora cervicornis*, to acute thermal stress. Using the Coral Bleaching Automated Stress System (CBASS)^39,40^, we analyzed the AS landscape of corals exposed to three levels of increasing temperature stress–Low (33℃), Medium (35℃), and High (37℃). Importantly, longitudinal sampling across six timepoints over a 19-hour time series captured conserved and dynamic shifts in the splicing profiles of the corals during heat exposure, as well as during a recovery phase subsequent to the thermal stress challenge. Our findings show evidence that corals may have evolved to utilize AS as a reversible mechanism for thermal acclimation to low levels of heat stress, but at higher levels of heat stress AS becomes irreversible and is associated with decreased photosynthetic efficiency of the symbionts (*Symbiodium ‘fitti*’^41^, the species known to associate with *A. cervicornis*^42^).

## Methods

### Acute thermal-stress assays

In June 2021, 22 coral fragments from each of eight *A. cervicornis* genets (defined as a genetically unique colony or group of colonies), sourced from the Mote Marine Laboratory *in situ* nursery (average 6.5 m depth, maximum depth 7.8 m), were evaluated for their thermal tolerance using the Coral Bleaching Automated Stress System (CBASS)^39,40^. Genets included Mote genet IDs 07, 31, 34, 41, 48, 50, 62, and CM5. The experimental system comprised four 20 L flow-through tanks fed from a common source water tank. Each flow-through treatment tank had an independent temperature profile controlled via an Arduino Mega 2560-based custom controller^40^. The system received an approximate flow-through rate of ∼1.5 L h^-1^ of fresh seawater collected at the nursery site and seawater supplied from the Mote Marine Laboratory water system at a ratio of ∼4:1. Water parameters of the source water, including average seawater temperature (29.1C +/- 0.82 Stdev, *n* =9), dissolved oxygen (83.2% +/- 2.18 or approximately 5.21 mg/L +/- 0.25), salinity (37.80 psu +/- 0.75) and pH (8.04 +/- 0.02) were monitored with a YSI Handheld Meter (YSI Yellow Springs, Ohio). The four treatment tanks’ temperature profiles included an ambient 30°C profile, a baseline control based on the local maximum monthly mean temperature for the area (MMM), and three heat stress treatments at 33°C (Low, +3°C MMM), 35°C (Medium, +5°C MMM), and 37°C (High, +7°C MMM) (Fig. 1a). An initial experiment had been performed with temperature set points of MMM (30°C), 34°C, 37°C, and 39°C, but nearly complete mortality was observed in the 37°C and 39°C treatments, necessitating a repeat of the experiment with reduced set points.

**Fig. 1:**
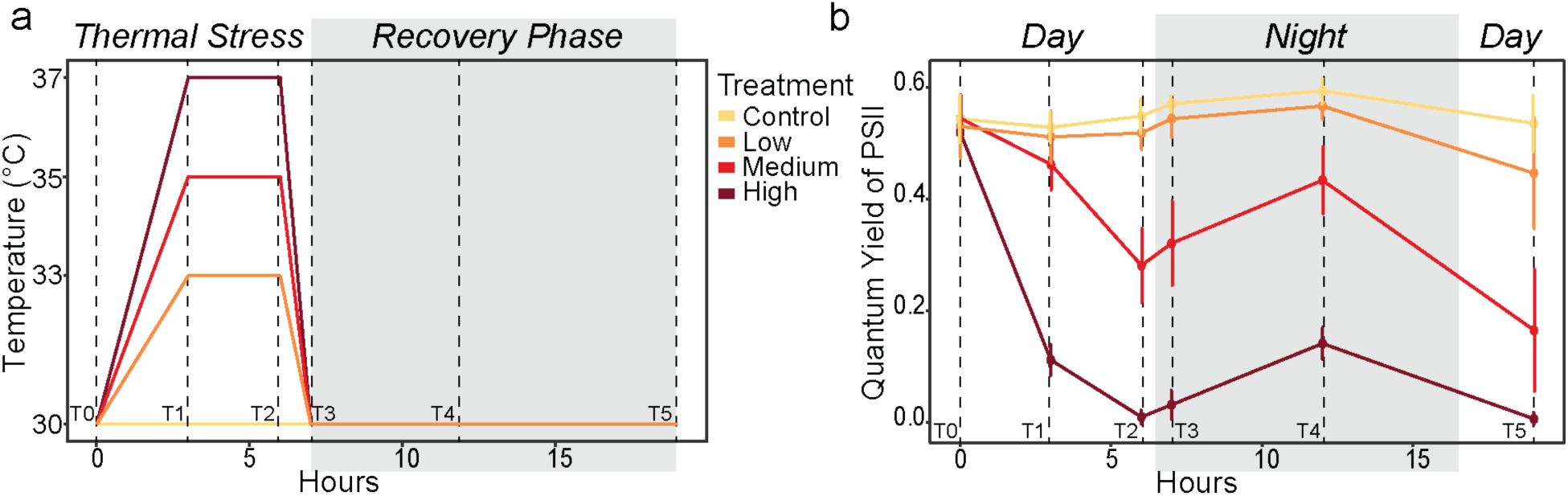
Acute thermal stress assay experiment design and phenotypic consequence on quantum yield of PSII. **a)** CBASS temperature profile design over the 19 hr assay (modified from Voolstra et al. 2020) run at Maximum Monthly Mean (MMM) at site (30°C; Control), 3 hr ramp- up followed by 3 hr heat-hold at MMM+3°C (Low), MMM+5°C (Medium), MMM+7°C (High), 1 hr ramp-down to 30°C (Control) temperature, and a 12 hr overnight recovery. RNASeq sampling timepoints are labeled T0-T5 with dashed lines. **b)** Variation in the maximum (dark-adapted Fv/Fm, night time) and effective (light-adapted ΔF/Fm’, day time) quantum yield of PSII at each timepoint. Error bars represent standard deviation of the mean. Pulse-amplitude modulated fluorometer measurement timepoints are labeled T0-T5 with dashed lines.

The assay was conducted over 19 hours, starting at 14:00 (T0), followed by a 3-hour temperature ramp-up (T1), a 3-hour hold at the respective treatment temperatures (T2), a 1-hour ramp-down to ambient (Control) temperature (T3), and a 12-hour overnight reprieve (T4 at 02:00+1, T5 at 09:00+1). The Control temperature tank remained unaltered throughout the experiment, while ambient light in all four tanks (white/blue LED artificial light at 600 µmol quanta m⁻² s⁻¹) was controlled in a fixed square-pulse photoperiod of light-on at 14:00, light-off at 20:30 after heat- load and light-on at 06:30+1. Samples were taken from each genet immediately after returning to shore (TF, Field) and after ∼ 1.5 h acclimation to the CBASS systems (T0, Initial) with the goal of distinguishing handling and tank effects from experimental effects. A total of 176 fragments were sampled for RNAseq - eight (one per genet) for Field samples (TF), eight for Initial (T0) samples, and 8 × 4 temperatures (Control, Low, Medium, and High treatments) × 5 timepoints (T1, T2, T3, T4, and T5). Coral fragments from all eight genets were evenly distributed across the treatment tanks and each genet was evaluated at each timepoint. One fragment from each genet was sampled at each time point in each treatment and immediately preserved in RNAlater (Thermo Fisher Scientific, USA). Samples were stored at 4C overnight to allow the buffer to penetrate the sample before being moved to -20C.

### Physiological measurements

Corals exposed to heat stress were analyzed by measuring the temperature-dependent loss of photosystem II (PSII) function, quantified through Quantum Yield of PSII. Specifically, effective photochemical efficiency (light-adapted ΔF/Fm’) was measured at timepoints T1, T2, and T5, while maximum photochemical efficiency (dark-adapted Fv/Fm) was measured at T3 and T4 (Fig. 1). Measurements were taken in the center of the coral fragment with a pulse-amplitude modulated fluorometer (Diving-PAM II; Heinz Walz, Effeltrich), setting the distance between sample and fiberoptics at approximately 10 mm, such that a sample at standard settings give a signal of 200- 400 units (basal fluorescence) throughout the experiment. Although temperature response curves have been commonly performed in the CBASS for Effective Dose temperature tolerance determinations^43^, here we focused on the dynamic change of Quantum Yield of PSII to compare physiological responses with alternative splicing dynamics across timepoints and temperature treatments. The statistical analyses and data plotting were conducted in R v.4.3.2 (R Core Team 2018)^44^.

### RNA extraction from coral tissue

RNA from 176 samples of *A. cervicornis* was extracted using a modified protocol of the QIAGEN RNeasy kit which utilizes TRIzol (Invitrogen, Waltham, MA) and chloroform to separate nucleic acids from proteins prior to binding nucleic acids to spin columns (as in https://openwetware.org/wiki/Haynes:TRIzol_RNeasy)^45^. For each extraction, tissue from 3-5 polyps was cut away from RNALater preserved coral fragments to avoid the inclusion of excess skeletal material. Column-bound nucleic acids were treated with DNAse (QIAGEN, Germantown, MD) to remove residual DNA. Concentrations and purity were measured by NanoDrop ND-1000 (ThermoScientific). RNA isolations were considered for further purification if they had an A280 ratio under 1.5, indicating potential contamination from TRIzol. RNA extractions sent for sequencing ranged from concentrations of 0.37 - 719.02 ng/µl with a median concentration of 154.35 ng/µl. Across all samples, RNA integrity (RIN) had a median value of 6.6. These extractions were sent for library preparation and sequencing on an Illumina NovaSeq 6000 S4 at Novogene.

### RNA-Seq read processing and alignment

Quality control of raw reads were processed and subsequently aligned using a custom RNA-Seq analysis pipeline (v0.2.7 available at https://github.com/baliga-lab/Global_Search). Briefly, reads were trimmed to remove adapter sequences and low quality reads using the TrimGalore v0.6.7 wrapper for Cutadapt v4.2 under default settings^46^. Trimmed reads were aligned to a concatenated host (*Acropora cervicornis*^47^) and symbiont (*S. ‘fitti’*^41^) metagenome using STAR v2.7.10a^48^ in twoPass Basic mode with filtering flags --outFilterMismatchNmax 10 -- outFilterMismatchNoverLmax 0.3 --outFilterScoreMinOverLread 0.66 --outFilterMatchNmin 0. The genets that yielded the host genome for *A. cervicornis* and the symbiont genome for *S. ‘fitti’* also stemmed from the Florida Reef Tract.

### Differential gene expression analysis

Following RNA-Seq alignment, aligned reads were split into symbiont (*S. ‘fitti’*) and coral host (*A. cervicornis*). Next, transcript abundances were determined with htseq-count v2.0.2^49^. Counts files were imported into R and differential gene expression analysis was conducted using the R package ‘DEseq2’ v1.38.3^50^. Transcripts with low read counts (<10) were filtered and expression analysis was conducted using the “DESeq” function. The lfcShrink function for shrinking log_2_ fold change estimates was applied to account for large dispersion with low read counts. Differential expression analysis was conducted for each heat stress treatment (Low, Medium, High) at each timepoint (T1-T5) against the Control at the same timepoint. Transcripts were considered significantly differentially expressed if s-values^51^ were smaller than 0.005 and were filtered for expression difference (log_2_ fold change > 1). Gene Ontology (GO) functional enrichment analysis^52,53^ was conducted using the R package ‘clusterProfiler’ v4.6.2^54^.

### Alternative splicing event detection and quantification

Alternative splicing (AS) classification and quantification was conducted using SplAdder v3.03^55^. RNA-Seq alignments were first split into species-specific datasets, i.e., host and symbiont and SplAdder analysis was conducted separately for each species. Briefly, splicing graphs were first generated for each sample separately and subsequently merged into a single joint splicing graph. During splice graph construction and augmentation, alignments were filtered according to the highest confidence level (--confidence 3) with five iterations to insert new introns. Graph nodes and edges were quantified on the merged graph of all samples. AS events were extracted using default SplAdder settings (allowing a maximum of 500 edges in the graph). The relative prevalence of a splice event was quantified using the widely used percent spliced-in (PSI) metric. PSI is the ratio of sequencing reads supporting an event type (e.g., exon skip) over the number of reads supporting alternative outcomes^56^.

Called AS events were analyzed using a workflow implemented as custom R scripts. Events with PSI values greater than 0.5 (i.e., majority usage) were tabulated per gene per sample and summarized by event type (alternative 3’ splice site, alternative 5’ splice site, exon skip, intron retention, multiple exon skips, and mutually exclusive exons), heat treatment (Field, Initial, Control (30°C), Low (33°C), Medium (35℃), and High (37℃)), and timepoint (T1 – T5). For global analysis, samples with high proportions of missing event calls (> 60%) due to insufficient read alignment coverage were filtered from downstream analysis. Finally, any events with missing data across the remaining samples were removed. This constrained set of AS events was used for principal component analysis (PCA), as implemented in the ‘prcomp’ function of the R package ‘stats’ (Base R v 4.2.1). The distribution of PSI estimates by temperature treatment and timepoint was examined, however, for visualization purposes, only events with the highest variance across samples (var > 0.04, *n* = 126) were plotted. To evaluate replicate agreement, variance between replicate genets for each AS event for each combination of timepoint and treatment was calculated. Further, to account for differences in the number of replicates for each treatment-timepoint combination due to quality filtering, we downsampled to a random selection of four genets such that variance was calculated on the same number of replicates for all sample types.

### Differential splicing analysis

Differential splicing analysis was conducted using the ‘test mode’ of the SplAdder software under default parameters for each combination of temperature (Low, Medium, High) and timepoint (T1- T5) against the Control at the same timepoint. This method models read counts at junctions using a negative binomial distribution with a generalized linear model for testing between groups and a multiple testing correction method of Benjamini-Hochberg^57^. Testing was conducted for each event separately. Prior to testing, samples with few called events due to poor read coverage were removed from testing groups (*n=*19 total across all tests). Delta PSI values (the difference between the mean-PSI values of the two groups being compared) were filtered for the most highly divergent events (delta PSI > 0.3^58,59^). Significant events (adjusted *p*-value < 0.05) were used in a Gene Ontology (GO) functional enrichment analysis^52,53^ with the R package ‘clusterProfiler’ v4.6.2^54^.

## Results

### Physiological response

To characterize the AS landscape of *A. cervicornis*, we evaluated the thermal response of 22 fragments from each of eight genets that were subjected to acute heat-stress profiles using the CBASS protocol. Measured tank temperatures closely reflected temperature profile set points (Supplementary Table 1). Heat stress was successfully induced as indicated by changes in Quantum Yield of PSII and fragments visually bleached in the Medium and High temperature treatments (Fig. 1b). Heat stress led to significant reductions in effective photochemical efficiency (light adapted ΔF/Fm’) after three hours of heat exposure (T2). This photodamage was more pronounced in the Medium and High temperature treatments, resulting in incomplete recovery following the stress period (T3 and T4; dark adapted). Even after the overnight recovery phase, corals did not fully regain PSII activity by T5 (light adapted; Fig. 1b).

### The global landscape of alternative splicing in *Acropora cervicornis*

We detected splice events using splice graphs generated from all samples across all conditions: Field, Initial, Control (30℃), Low (33℃), Medium (35℃), and High (37℃) at all timepoints (Fig. 1b). Fewer mapped reads for the symbiont (on average, 6.7-fold fewer than the host; Supplementary Table 2) resulted in high proportions of missing event calls across samples and low AS event counts relative to the host (Supplementary Fig. 1). This proved insufficient for differential AS testing between treatment groups and, thus, all subsequent analyses focus solely on the coral host, *A. cervicornis*. Further, no samples for High treatment at timepoint T5 passed the filtering thresholds resulting in 156 samples for global analysis. After all filtering was complete, 156 samples total passed the quality thresholds for global analysis. The SplAdder software detected and quantified six classes of splice events: alternative 5’ or 3’ splice sites, exon skips, intron retention, multiple exon skips, and mutually exclusive exons. In total, we characterized 137,799 splice events in 11,984 genes, comprising 40% of the *A. cervicornis* genome (Table 1; Supplementary Table 3). The majority (69%) of AS events detected were actively used during the experiment (defined as having at least one sample with PSI >= 0.5; Table 1). Across all samples (each replicate at each timepoint for each temperature treatment), alternative 3’ splice sites and intron retention events were most prevalent, while multiple exon skips were less common (Fig. 2a). The splicing profile was stable (identical PSI values) across all samples in 11% of events. We found a high level of concordance across replicate samples with overall low variance in PSI estimates between replicates (Supplementary Fig. 2) and similar splicing profiles for each of the eight genets for each combination of treatment and timepoint (Fig. 2b). Together, the most conservative settings for calling AS events, subsequent downstream quality filtering, and the reproducibility of findings across replicates indicates that the splicing estimates and event types are not artifacts and can be confidently linked to mechanisms associated with biological response.

**Fig. 2:**
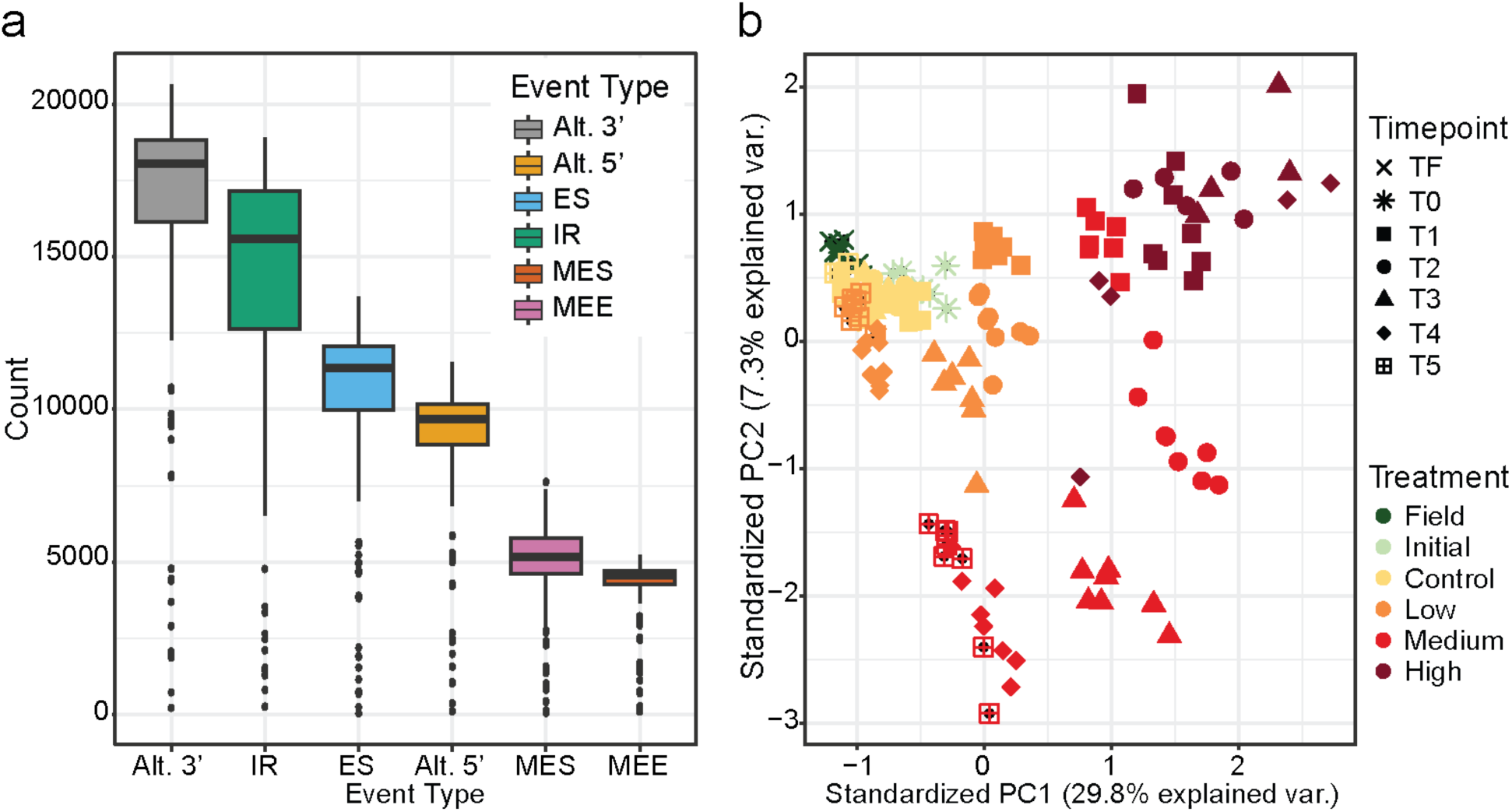
Summary of all AS events across samples. **a)** The distribution of counts of AS events (PSI > 0.5 in at least one sample) by event class (gray = alternative 3’ splice site (Alt. 3’), yellow = alternative 5’ splice site (Alt. 5’), blue = exon skip (ES), green = intron retention (IR), red = multiple exon skips (MES), pink = mutually exclusive exons (MEE)). **b)** Principal Component Analysis (PCA) of AS events for all samples included in global analysis (*n* = 156) with colors indicating treatment (Field = dark green, Initial = light green, Control = yellow, Low= orange, Medium = red, High = dark red) and shapes indicating timepoints (TF (Field) = cross ; T0 (Initial) = star; treatment timepoints: T1 = square, T2 = circle, T3 = triangle; recovery timepoints: T4 = diamond, T5 = square plus).

**Table 1:** Summary statistics of AS across all *Acropora cervicornis* samples. The number of alternatively spliced genes and number of splice events per event type is shown for actively used (at least one sample with PSI >= 0.5) and all detected AS events in all samples (*n* = 176).

| Event type | Number of genes (Number of events) |  |
| --- | --- | --- |
|  | Active usage | All |
| *Alt 3' splice site* | 8,022 (23,375) | 8,914 (33,468) |
| *Alt 5' splice site* | 6,269 (14,136) | 8,056 (24,983) |
| *Exon skip* | 6,723 (16,456) | 8,231 (34,551) |
| *Intron retention* | 6,127 (24,262) | 6,366 (25,894) |
| *Multiple exon skips* | 2,980 (5,883) | 3,026 (6,174) |
| *Mutually exclusive exons* | 1,567 (11,056) | 1,808 (12,729) |
| **Total** | 11,350 (95,168) | 11,984 (137,799) |

### The alternative splicing response of the coral host to thermal stress

In addition to characterizing the composite splice profile of *A. cervicornis*, we examined cases where AS events differed among temperature treatments and timepoints. Overall, the splicing response varied based on the level of heat stress, as shown by the clustering of colonies by temperature (Fig. 2b). We also observed separation of samples by timepoint with an increasing spread between temperature treatment (T1-T3) and recovery phase timepoints (T4 and T5) with increasing levels of heat. The effect of the CBASS tank was minimal, as indicated by the tight clustering of Field, Initial, and Control samples (Fig. 2b; see also Supplementary Fig. 3). Additionally, Low level thermal stress (33°C) showed less pronounced separation from Control conditions, most notably during recovery phase timepoints, suggesting recovery indeed occurred with respect to AS. All six classes of splice events (alt. 3’ splice site, alt. 5’ splice site, intron retention, exon skipping, multi-exon skipping, and mutually exclusive exons) were represented in the top most variable events, with intron retention dominating (Fig. 3a). These events show distinct patterning in PSI by thermal stress with higher PSI values during higher temperature treatments and lower PSI values during Control conditions as well as recovery phase timepoints following Low heat treatment (Fig. 3).

**Fig. 3:**
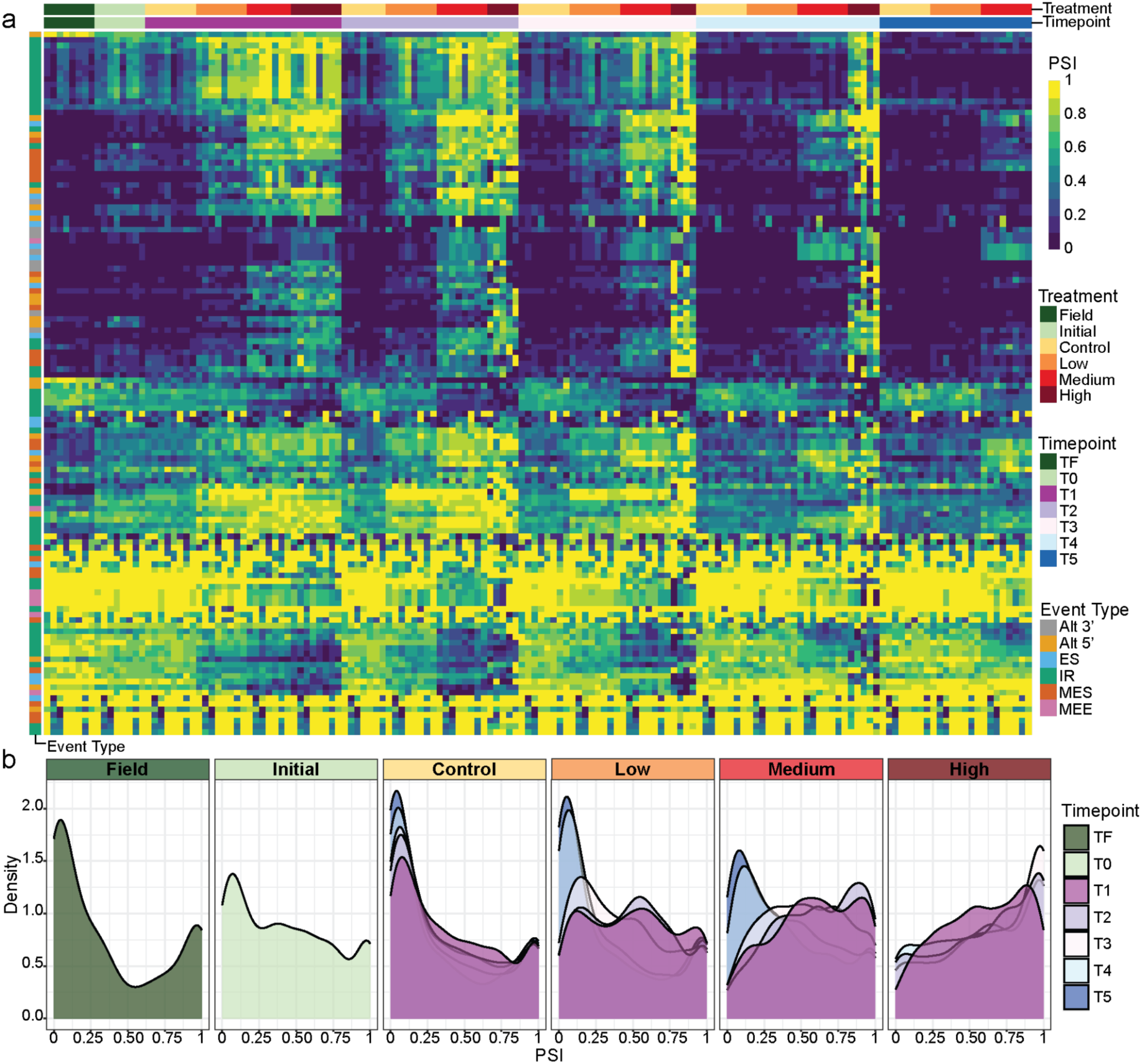
**Thermal stress and recovery phase-associated dynamic changes in Percent Spliced In (PSI) for global AS events with the highest variance**. The PSI (ranging from 0 - 1) of the top (var > 0.04; *n* = 126) most variable events across all samples included in global analysis (*n* = 156) is indicated by **a)** a heatmap with rows clustered by euclidean distance (NA values indicated by gray) and **b)** density distributions for each treatment type (Field = dark green, Initial = light green, Control = yellow, Low= orange, Medium = red, High = dark red) at each timepoint (TF (Field) = dark green; T0 (Initial) = light green; treatment timepoints: T1 = magenta, T2 = light purple, T3 = light pink; recovery phase timepoints: T4 = light blue, T5 = dark blue). Splice events are labeled along the left y-axis (gray = alternative 3’ splice site (Alt. 3’), yellow = alternative 5’ splice site (Alt. 5’), blue = exon skip (ES), green = intron retention (IR), red = multiple exon skips (MES), pink = mutually exclusive exons (MEE)). Rows are clustered using the default clustering method “complete” in ‘pheatmap’ v1.0.12.

To elucidate the influence of heat stress (treatment) over time (timepoint) on the complexity of AS, we examined the distribution of PSI values across splice events with the highest variance across all samples and observed a slight increase in the PSI distribution of Initial samples relative to Field samples. Shifts in the distribution of PSI values also indicate increased AS with higher heat stress treatment. For each heat treatment (Low, Medium, and High), there was a shift in the distribution of PSI values, which indicated more AS during heat treatment (T1-T3) relative to recovery phase (T4 and T5; Fig. 3b). Under the Low heat treatment (33℃) there was a return to baseline levels of AS (i.e., similar to Control (30℃)) during the recovery phase timepoints. This shift was also observed under Medium (35°C) heat treatment, albeit to a lesser degree with some persistence in increased AS. Increased AS persisted at elevated levels even during the recovery from High (37℃) temperature treatment (Fig. 3b).

### Differentially spliced genes are dominated by intron retention

We investigated the differential AS of events across pairwise contrasts of sample groupings for all heat treatments at all timepoints. Intron retention events were the most abundant class of significant AS events relative to Control conditions (49% of all significant events across all tested contrasts; Fig. 4a). Few significant events were detected between Low and Control treatments (*n* = 261 total across all timepoints; *adj. p* > 0.05; delta PSI > 0.3), whereas High (*n* = 4,600 total across all timepoints; *adj. p* > 0.05; delta PSI > 0.3) and Medium (*n* = 4,402 total across all timepoints; *adj.p* > 0.05; delta PSI > 0.3) heat treatments show many cases of significant differential AS events (Supplementary Table 4). To account for the effect of sampling and transition of colonies to the CBASS, we also tested for differential AS between Field, Initial, and Control conditions and found few significant differences (*n* = 1,044 total across all eleven contrasts; *adj. p* > 0.05; delta PSI > 0.3) in AS across each of these contrasts (Supplementary Fig. 3). Additionally, results indicate a greater number of significant differential AS during heat treatment timepoints (T1-T3) relative to recovery phase timepoints (T4 and T5). Genes involved in significant differential AS relative to Control conditions were enriched for several cellular functions, including heat shock transcription factors, splicing machinery, and splicing regulation itself (Fig. 4b; Supplementary Table 5).

**Fig. 4:**
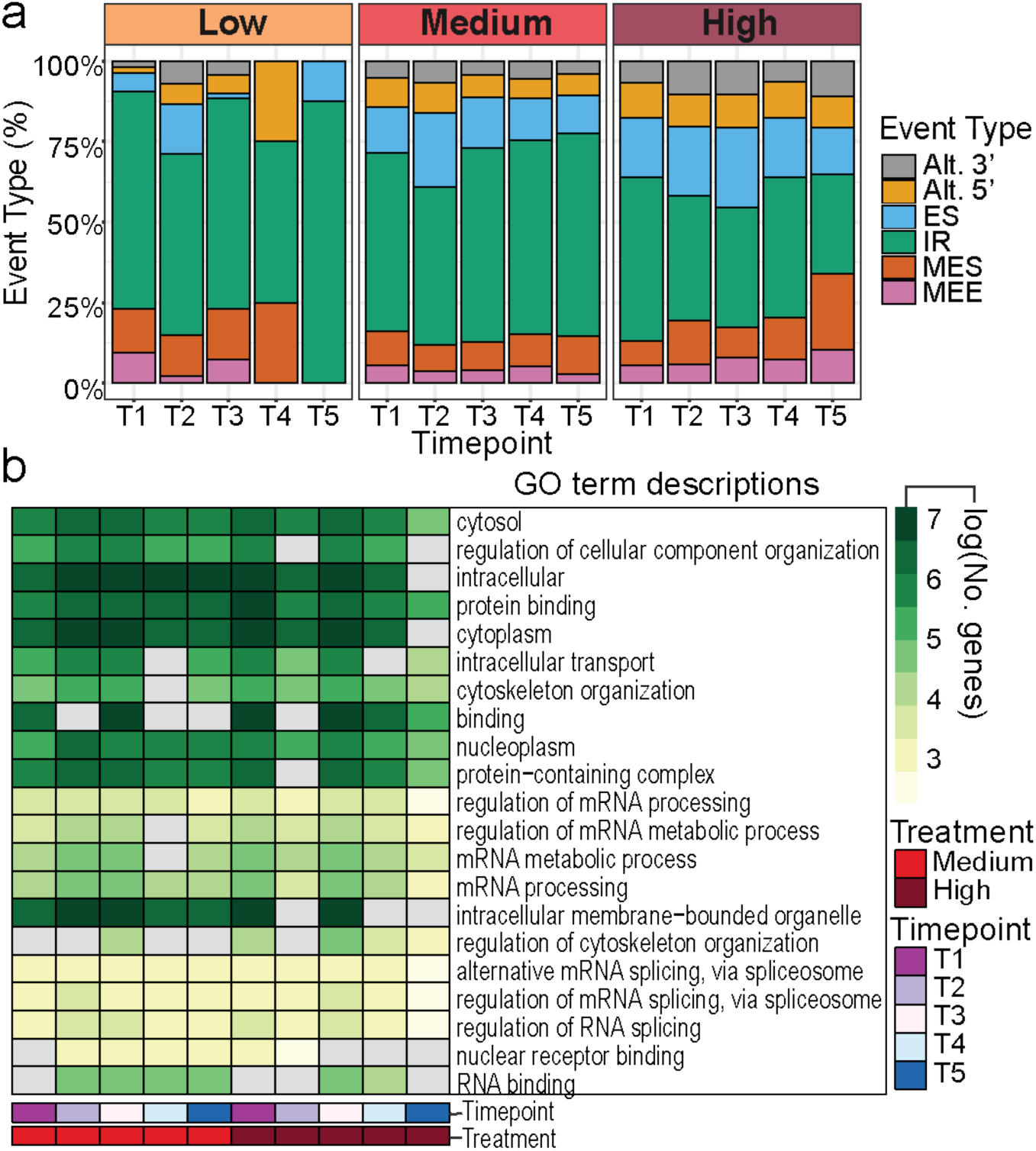
Proportions of significantly differential AS events and distributions of AS genes across biological processes. **a)** The percent of significantly differentially alternatively spliced events (*adj. p-value* < 0.05; delta PSI > 0.3) for each heat treatment Low (33℃), Medium (35℃), High (37℃) at each timepoint (T1 - T5) against Control (30℃) conditions is binned according to the splice event type (gray = alternative 3’ splice site (Alt. 3’), yellow = alternative 5’ splice site (Alt. 5’), blue = exon skip (ES), green = intron retention (IR), red = multiple exon skips (MES), pink = mutually exclusive exons (MEE)). **b)** Heatmap of the log of the gene count (NA values indicated by grey) for the top five most significantly enriched GO terms (labeled by GO term description) for each timepoint (T1 = magenta, T2 = light purple, T3 = light pink, T4 = light blue, T5 = dark blue) for Medium (35℃; red) and High (37℃; dark red) temperature treatments relative to Control (30℃).

### Overlap of differential AS with differential gene expression

We assessed the degree of overlap between differential gene expression and AS across all treatment vs Control comparisons. At Medium and High levels of heat treatment, across all timepoints, between 350 (33%) and 1,109 (52%) significantly differentially AS genes (relative to Control) were also significantly differentially expressed (Fig. 5a). At Low heat treatment, between 0 and 82 (16%) differentially AS genes were also differentially expressed. For all treatments, the highest degree of overlap occurred during treatment (T2 and T3) with a decrease in overlap during the recovery phase (T4 and T5). Additionally, we observed that many of the same AS events were maintained across treatments (i.e., the same intron is retained; Fig. 5c; see also Supplementary Table 4). Differential gene expression analysis showed disproportional downregulation of genes relative to upregulation across all treatments and timepoints (Supplementary Fig. 4a). However, of the genes that were differentially AS (*n* = 2,550), Low at T1, Medium at T1, and High at T2-T5 had a higher proportion of upregulated genes than downregulated genes.

**Fig. 5:**
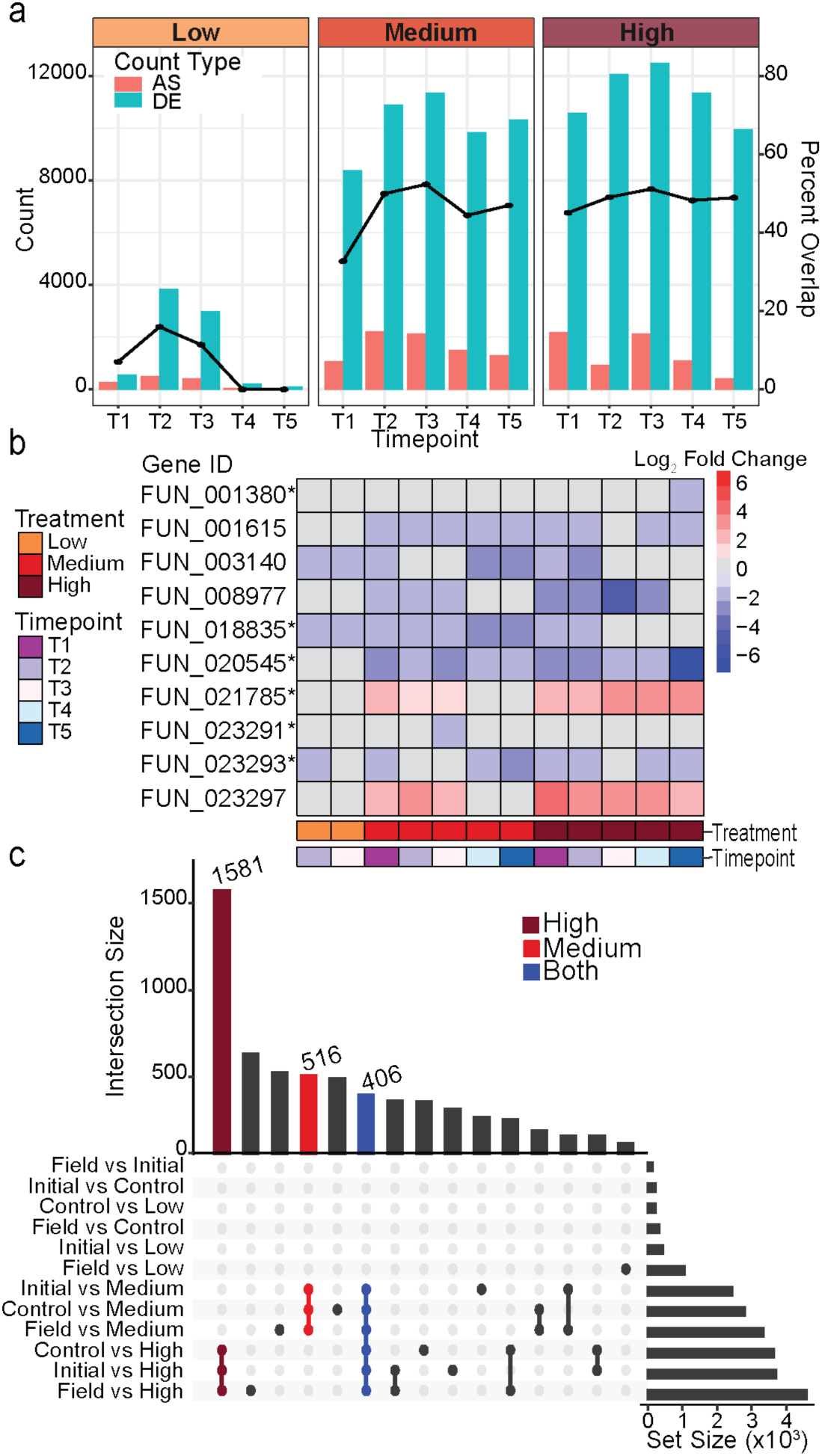
Comparison of genes associated with differential expression and AS events during thermal stress and recovery phase. **a)** Percent overlap (right y-axis) of genes that are significantly differentially AS with genes that are significantly differentially expressed (DE) is shown as a line graph for each treatment (Low, Medium, and High) at each timepoint (x-axis; T1- T5) relative to Control at the same timepoint. Bars display the total count (left y-axis) of genes with significant AS (pink) or significant DE (blue) for each contrast. **b)** Log_2_ fold change values for the gene expression of the ten genes with putative NMD-related functions are shown for each temperature treatment (Low, Medium, High) at each timepoint (T1-T5) relative to Control conditions at that same timepoint. Results are shown if at least one gene was significantly DE for the given comparison. Positive log_2_ fold change values are shown in red hues while negative log_2_ fold change values are shown in blue hues. Grey indicates no significant DE. Gene IDs are denoted with an (*) if the gene was significantly differentially spliced in at least one comparison. **c)** UpSet plot showing the intersection of significant alternative splicing events for each temperature treatment contrast (time points aggregated) with shared events indicated in dark red (all High contrasts), red (all Medium contrasts), and blue (all High and Medium contrasts).

Across all temperature treatments, upregulated genes were enriched for GO terms related to heat stress response (e.g., regulation of cellular response to heat, response to heat, response to temperature stimulus, heat shock protein binding), protein refolding (chaperone cofactor-dependent protein refolding), and regulation of transcription (regulation of transcription by RNA polymerase II, regulation of DNA-templated transcription, and negative regulation of transcription by RNA polymerase II; Supplementary Table 6). In contrast, downregulated genes were enriched for GO terms that included DNA repair and replication, and many involving metabolic processes (e.g., small molecule metabolic process, organic acid metabolic process, and carboxylic acid metabolic process; Supplementary Table 6). Together these findings suggest that during heat stress the coral downregulated DNA repair and metabolism while upregulating mechanisms for heat stress management.

### Expression patterns of the nonsense mediated decay (NMD) pathway

Because studies have suggested that AS may fine-tune gene expression through coordinated coupling of AS with nonsense mediated decay (NMD), a cellular surveillance mechanism that detects and degrades mRNA transcripts containing premature stop codons^60,61^, we examined the AS and expression patterns of NMD-related genes. Altogether, of the ten genes with putative functions in the NMD pathway, six underwent significant differential AS (Fig. 5b). Relative to Control conditions, across all temperatures and timepoints, the majority of these genes underwent either no significant change in gene expression or were downregulated (eight out of ten genes; Fig. 5b). Notably, only two NMD-related genes were upregulated (gene IDs FUN_023297 and FUN_021785) during Medium (timepoints T1-T3) and High (all timepoints) heat treatments (Fig. 5b).

## Discussion

This study represents the first systems level characterization of the transcriptome-wide landscape of AS in a coral experiencing and recovering from heat stress. To our knowledge, only two previous studies have explicitly examined AS in corals. Huang et al.^62^ detected and quantified five modes of AS in response to symbiont infection and found that intron retention predominated in *Acropora digitifera*. Another study investigated gene duplication and differences in cDNA length (which they attribute to AS) specifically in a gene from the fluorescent protein gene family in a single individual of *Montipora spp.*^63^. As early branching metazoans, corals represent an important taxonomic group for studying AS due to the documented differences between animals and plants^64–66^. Though corals are animals, we found that the stress-responsive AS profiles resemble that of many plants. While exon skipping is the most common class of AS observed in animals, our finding of alternative 3’ splice sites and retained introns in *A. cervicornis* is more reminiscent of splicing patterns observed in plants^64,67–70^ (Fig. 2a; Fig. 4a; Supplementary Fig. 3). Consistent with this observation, the proportion of genes undergoing AS in *A. cervicornis* (∼40%; Table 1), while lower than that of higher order metazoans (∼60%), is higher than that of other non-chordates (∼26%) and more similar to that of plants (∼31%)^71^. This is particularly interesting as the degree of AS has been reported to be proportional to organismal complexity (as measured by the number of distinct cell types) in the metazoan lineage, with an independent rise in plants^71^. The only other cnidarian included in the study of AS vs. complexity, the sea anemone *Nematostella vectensis*, was also observed to have higher levels of AS (32%) than other non-chordate metazoans^71^. Considering the prominent role of gene family expansions in both evolution of multicellularity and AS, our findings may help investigate the divergent evolution and different roles of AS across metazoan lineages^72^.

Longitudinal sampling and multiple levels of heat treatment (Fig. 1a), allowed us to capture the influence of thermal stress, as well as immediate recovery dynamics in the coral host, *A. cervicornis* on alternative splicing, differential gene expression, and damage to PSII. Results showed the degree of AS increased proportionally with levels of acute thermal stress (Fig. 3). During recovery phase timepoints (T4 and T5), AS decreased (similar to Control, Field, and Initial conditions) following Low level heat stress, however, at higher levels of heat stress elevated AS persisted partially (Medium) or fully (High; Fig. 3b). Further, increase in AS corresponded to loss of photosynthetic efficiency of the algal endosymbiont (Fig. 1b), and was irreversible in the timeline of this experiment when heat stress exceeded a threshold, i.e., a tipping point, potentially due to lasting molecular damage. Determining such a threshold has become progressively more important as sea surface temperatures continue to rise and marine heatwaves become more frequent and intense due to climate change^73,74^.

Intron retention was the predominant type of differential AS event across treatments and time (Fig. 3a; Fig. 4a; Supplementary Table 4; Supplementary Fig. 3). Although commonly observed across many organisms and contexts^75^, the consequences of increased intron retention remain an open question^75^. Since intron retention is expected to lead to loss-of-function due to the introduction of premature termination codons (PTC), this AS event was historically assumed to be a consequence of a malfunction of the spliceosome. However, there is growing evidence that intron retention, and AS, in general, may represent a mechanism of post-transcriptional gene regulation^28,38,76–79^. In humans, where exon skipping is more common, intron retention has been tied to many diseases, including cancers and neurodegenerative diseases^80^, but in plants, intron retention is largely associated with the abiotic stress response^28,64–66^.

Corals have a biphasic life cycle with a dispersive larval stage and a sessile adult form, similar to plants. Consequently, like plants, adult corals must also acclimate to changing environmental conditions. Differential methylation, followed by differential gene expression is one-way corals and plants adjust their phenotypes to stress^27,81^. AS also plays an important role in mediating rapid response to heat stress in plants via modulation of mRNA abundance or subcellular localization, generation of functionally novel protein variants, altered domains or binding properties, production of regulatory long non-coding RNAs, allele-specific expression, and alteration of translation efficiency^82–85^. For example, in *Arabidopsis thaliana*, heat shock induces intron retention of heat shock transcription factorA2 (*HsfA2)*, producing a truncated protein which acts as a positive transcriptional activator to enhance the expression of *HsfA2* for heat tolerance^86^. Similarly, we discovered significant differential AS events in the transcript encoding Heat Shock Factor 1 (*HSF1*), during coral response to heat stress (Supplementary Table 4). It remains to be determined whether the AS transcript isoforms are destined for NMD, localized to the cytoplasm rather than exported to the nucleus, or part of a positive auto-regulatory feedback loop. However, given the recovery of the corals under Low heat treatments with respect to AS levels (Fig. 3b) and the documented reduction in thermal tolerance of corals following *HSF1* knockdown^87^, it seems likely that AS of this particular gene is important for heat stress response and further suggests that AS regulation may be an adaptation for sessile organisms to acclimate to environmental stressors.

The lower abundance of transcripts of eight of ten NMD-related genes suggested that AS transcript isoforms of other genes are potentially not degraded by this pathway. Two NMD-related genes that were upregulated include *SMG6* and *PABPC4* (FUN_021785 and FUN_023297, respectively; Fig. 5b). *SMG6* encodes a component of the telomerase ribonucleoprotein complex, which initiates NMD by acting as an endonuclease near PTCs^88,89^, and also plays a role in telomere maintenance^90^. On the other hand, *PABPC4* is an RNA-processing protein that binds to the 3’-poly(A) tail present in most eukaryotic mRNAs to effect negative regulation of NMD^91,92^. Therefore, increased expression of this gene is expected to suppress NMD. Taken together, these results imply that it is unlikely that transcripts with intron retention are degraded through the NMD pathway. Further, the NMD and unfolded protein response (UPR) pathways work in a reciprocal manner to coordinate an efficient response to stress, with the UPR suppressing NMD during endoplasmic reticulum stress and NMD suppressing the UPR during innocuous or low stress^93^. Indeed, genes encoding functions in the protein folding and refolding pathways were significantly upregulated during *A. cervicornis* response to heat stress (Supplementary Table 6).

With the multitude of mechanisms governing AS and its downstream consequences, splicing represents a complex regulatory layer with large gaps in our understanding of their impacts at the phenotypic level^28,94^. However, consistent with findings from this study, it is known that the regulators of AS are themselves often targets of AS during the thermal stress response across many organisms^94–100^. We observed significant differential AS of spliceosome components and associated splicing regulatory genes across all timepoints and heat treatments relative to Control conditions (Fig. 4b; Supplementary Table 5). Further, ∼50-70% of these genes were not differentially expressed under Medium and High temperatures (Fig. 5a). This suggests that AS may play a complementary, yet, distinct role in the thermal stress response that may not be captured by analysis of relative changes in expression of canonical transcript isoforms. For instance, *SF3B1*, a core subunit of the U2 snRNP complex of the spliceosome, regulates both the expression levels and activity of *HSF1* in *Caenorhabditis elegans* and humans^101^. Interestingly, *SF3B1* was differentially AS (but, notably, not differentially expressed) in *A. cervicornis*, whereas its target *HSF1* was both differentially AS and differentially expressed (Supplementary Table 4). This finding suggests that a similar mechanism of SF3B1-mediated regulation of HSF1 may be important for thermal stress response of corals.

The extent to which the global shifts in splicing in *A. cervicornis* represent splicing dysregulation, like in human disease^102^, or an adaptive response, as assumed in plants ^84^, requires additional study. In *A. thaliana*, ‘thermopriming’, i.e., exposure to sub-lethal heat stress prior to heat shock, induced a ‘splicing memory’ whereby non-primed plants showed increased intron retention during heat shock relative to primed plants^103,104^. The authors suggested that intron retention may represent a cellular reservoir of unspliced transcripts at the ready for flexible splicing upon heat stress relief. However, the decrease in intron retention in primed plants could also be interpreted as a correction for splicing dysregulation. Indeed, at the highest levels of heat treatment, wherein corals were unable to recover (Fig. 4b), there was greater variability between replicates (Fig. 2b; Supplementary Fig. 2), suggesting that in this context it may be the case that the pool of AS transcripts may have, at least partly, resulted from splicing machinery breakdown. Nonetheless, this is unlikely to be the case during response to Low heat stress, wherein patterns of AS had low variability across replicates. Further, our finding that the NMD pathway was largely suppressed during heat stress, indicates that AS transcripts, especially those with intron retention, are not marked for decay, and instead may encode adaptive functions. This conjecture is also supported by the observation that many of the same AS events were observed across treatments (Fig. 5c; Supplementary Table 4), which is suggestive of an actively regulated mechanism of targeted splicing, rather than a consequence of a breakdown of the splicing machinery. Finally, the finding that a significant number of splicing regulators were themselves targets for AS in *A. cervicornis* has also been reported in other organisms^94–100^, suggesting this mechanism of adaptive regulation may have emerged early in evolution, particularly in lineages of organisms with a sessile lifestyle. Although further studies are needed to delineate specific mechanisms and functions for AS, our findings demonstrate compelling evidence that this process and its products play an important role in mediating heat stress response in corals.

Here, we present the first evidence of global alternative splicing in response to acute thermal stress in a coral. The reproducibility and reversibility of AS patterns in Low heat stress, taken together with other observations (e.g., downregulation of NMD pathway genes) suggests that this process may play an important role in the thermal stress response of corals, similar to plants, which also have a sessile lifestyle. In that regard, the irreversibility and variability of AS in high levels of heat stress, suggests a breakdown of this process, which happens concomitantly with inability of the coral to recover even after relief from heat stress. The results presented here lay the foundation for future investigations to elucidate whether the breakdown of AS is a cause or consequence of the inability of coral to recover from heat stress. Further work examining whether AS also plays a role in the response of the algal endosymbiont, is consistent in other populations of *A. cervicornis* across the Caribbean, as well as conserved in phylogenetically divergent coral species, would clarify both the evolutionary history of AS in metazoans, as well as potential applications in conservation efforts.

## Acknowledgements

This work was supported by the Paul G. Allen Family Foundation (PGAFF). This is publication number 1 of the Global Search Consortium https://zenodo.org/records/7835728.

## Author contributions

JJV, NSB, DJB, CRV, IBB, LKB conceived the experiment, DJB, IBB, KEP, VZR, NSL, TLC performed the experiment, KEP, VZR, NSL, and TLC performed the laboratory analysis, KHS, JJV, NSB, DJB, CRV, IBB, NSL, KGC, ST, WW analyzed data, KHS, JJV, NSB, DJB, CRV, IBB, NSL, KGC, RA interpreted data, KHS led data analysis and wrote the manuscript with editing, input, and final approval from all authors.

## Data Availability Statement

Data reported in this publication is available under BioProject PRJNA1213837 (https://www.ncbi.nlm.nih.gov/bioproject/1213837) under the ‘Global Search for Genetic Regulators of Coral Resilience to Thermal Stress’ umbrella BioProject PRJNA749006 (https://www.ncbi.nlm.nih.gov/bioproject/PRJNA749006).

## Ethics declarations

### Competing interests

The authors declare no competing interests.

Supplementary Table 1: CBASS temperature profile log file

Supplementary Table 2: Count of uniquely mapped reads per sample per species (coral host and symbiont)

Supplementary Table 3: PSI estimates of called events for all samples in the coral host Supplementary Table 4: Significant differential AS events in the coral host

Supplementary Table 5: Enriched GO terms for significant differentially AS genes for each tested contrast

Supplementary Table 6: Enriched GO terms for significant differentially expressed genes for each tested contrast

---

**Supplementary Fig. 1:**
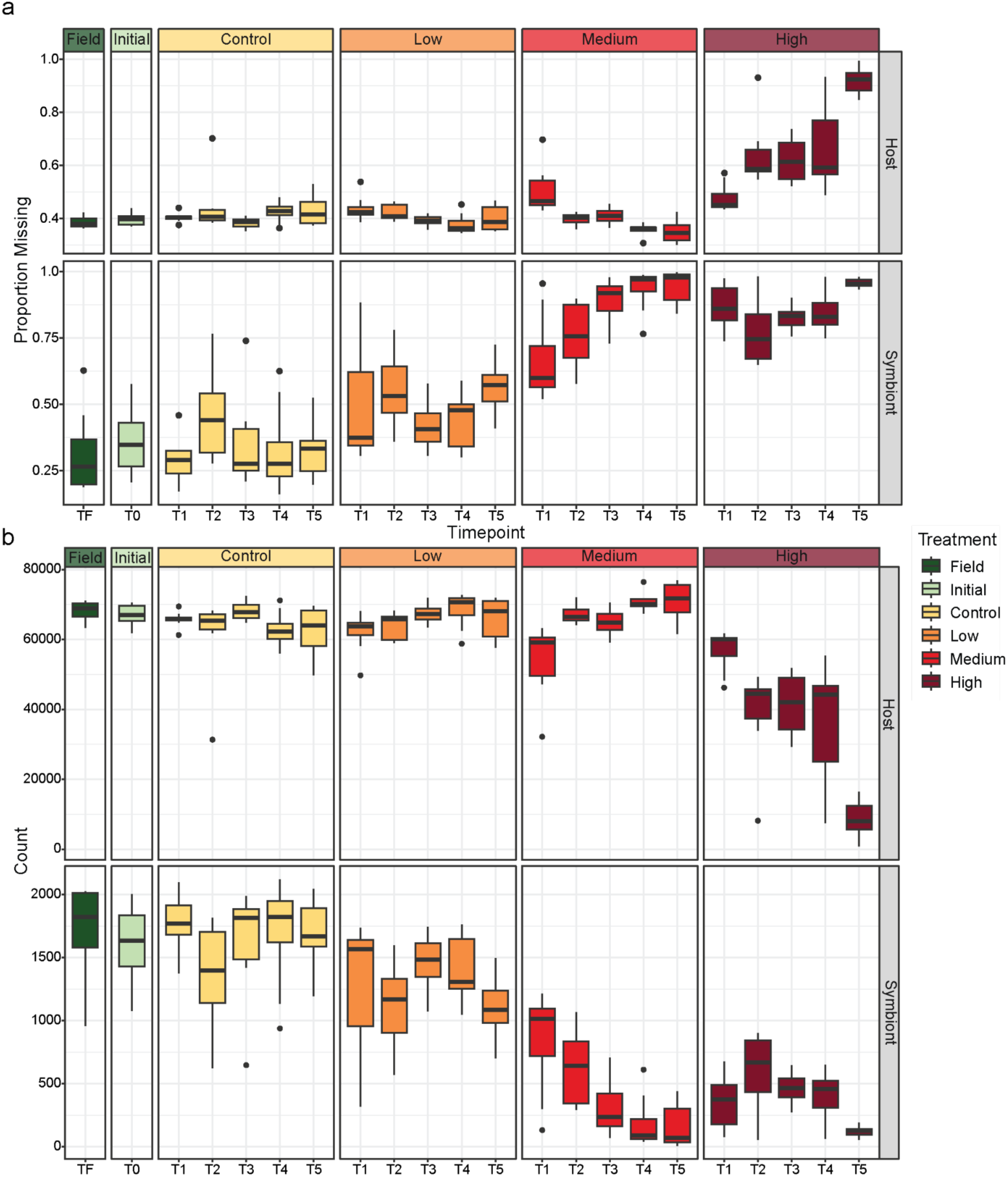
RNA-Seq and AS filtering quality control by species. Boxplots of **a)** the proportion of AS events with missing PSI estimates and, **b)** the total number of AS events are shown for each sample type (Field = dark green, Initial = light green, Control = yellow, Low= orange, Medium = red, High = dark red) at each timepoint (*x*-axis) for both the coral host and the algal endosymbiont. Note the different y-axis scales.

**Supplementary Fig. 2:**
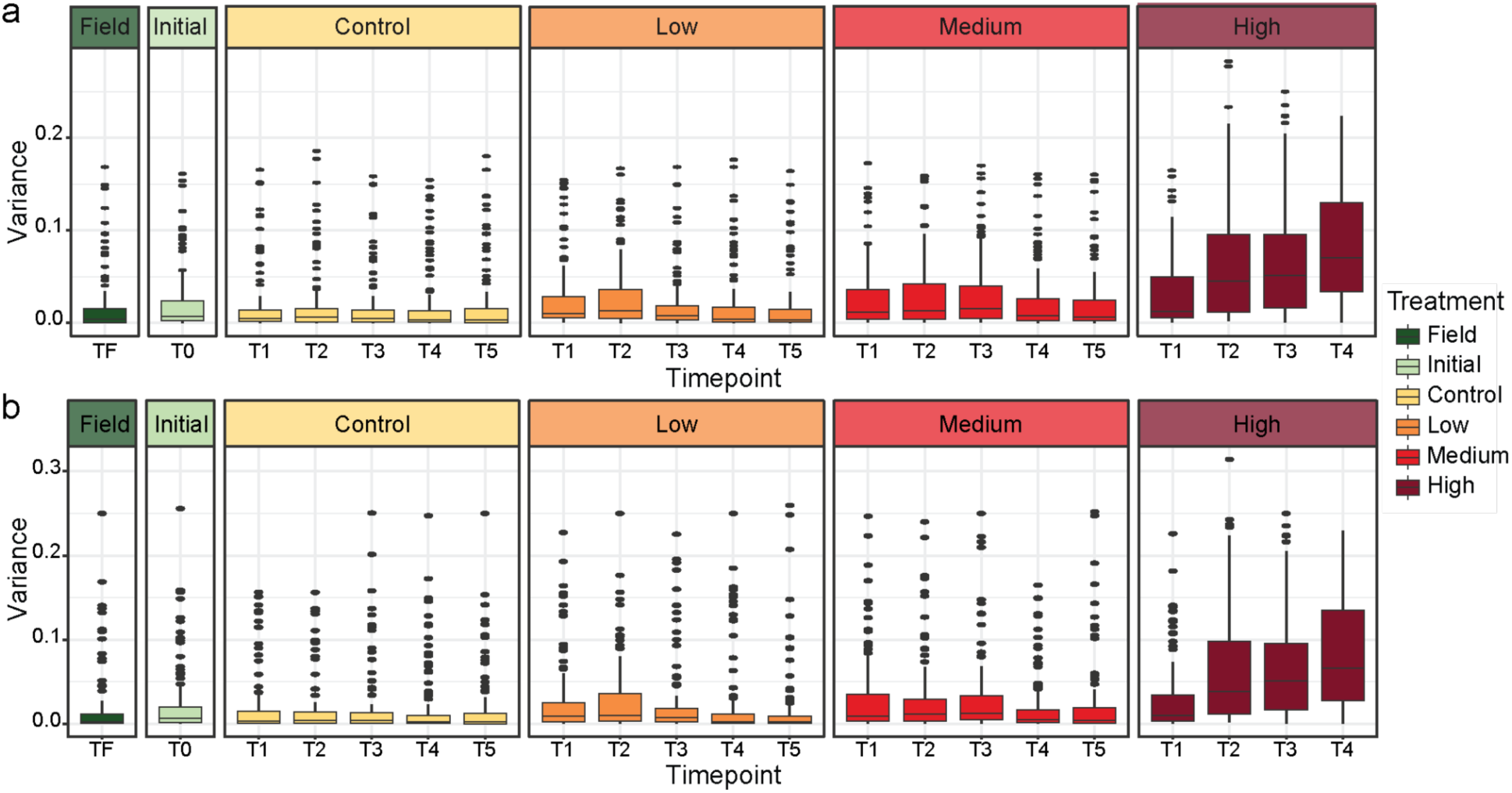
Within group variance by sample type in the coral host. Boxplots of the distribution of variances in PSI estimates per AS event by treatment type (Field = dark green, Initial = light green, Control = yellow, Low= orange, Medium = red, High = dark red) and timepoint (*x*-axis). Events with identical PSI for all samples or missing values were filtered prior to variance calculation. Plots are shown for **a)** all samples in the global analysis (*n* = 156) and **b)** four randomly selected genets for each sample category (*n* = 84).

**Supplementary Fig. 3:**
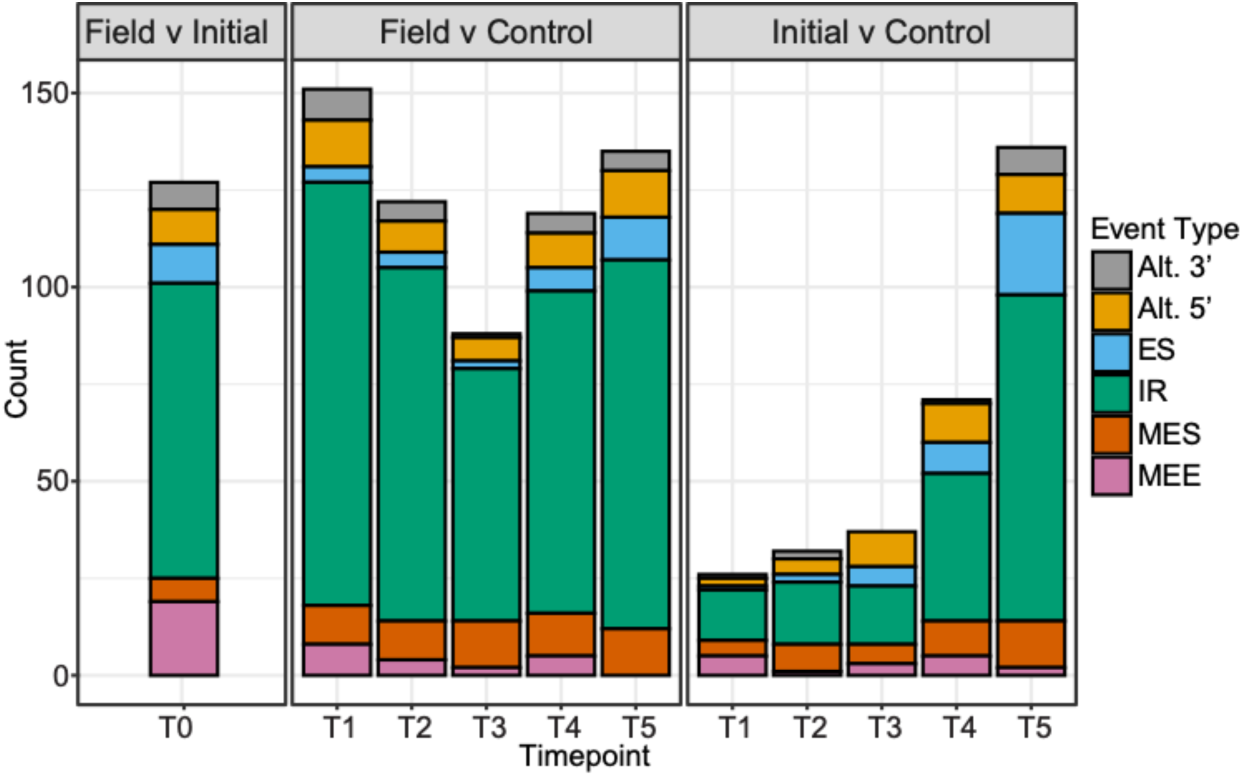
Significant differential AS in Field, Initial, and Control contrasts in the coral host. The number of significantly differentially alternatively spliced events (adj. p-value < 0.05; delta PSI > 0.3) for pairwise contrasts comparing Field, Initial, and Control samples at each timepoint. A count of the number of significant alternative splicing events for each contrast is binned according to the splice event type (gray = alternative 3’ splice site, yellow = alternative 5’ splice site, blue = exon skip, green = intron retention, red = multiple exon skips, pink = mutually exclusive exons).

**Supplementary Fig. 4:**
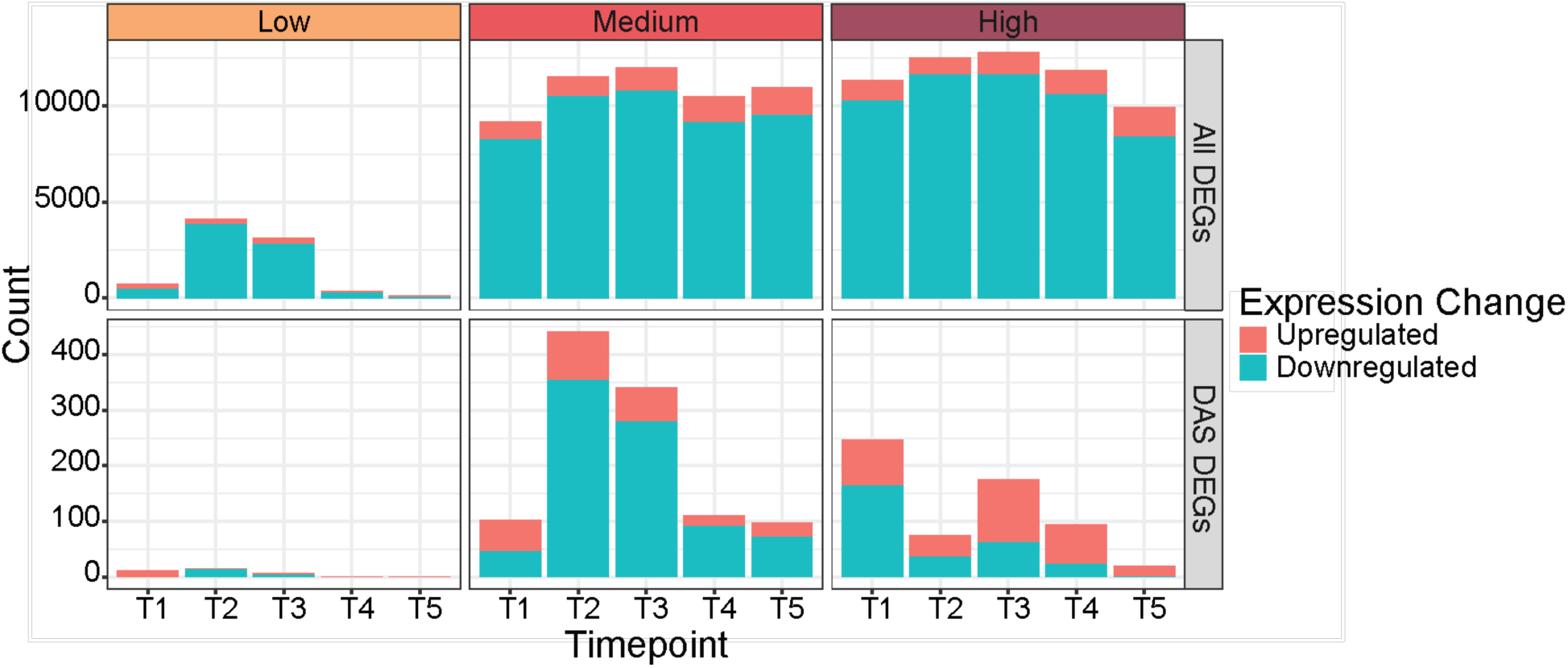
Summary counts of genes with differential expression by contrast. The number of significantly differentially expressed genes (DEGs) are shown for each temperature treatment (Low, Medium, and High) at each timepoint (T1-T5). Counts are shown for positive log_2_ fold change (upregulated; pink) and negative log2 fold change (downregulated; blue). The top panels show counts for all DEGs while the bottom panel shows counts only for DEGs that were also differentially alternatively spliced (DAS).

